# A Bayesian approach to interpret time-resolved experiments using molecular simulations

**DOI:** 10.1101/2025.07.19.665657

**Authors:** Carl G. Henning Hansen, Simone Orioli, Kresten Lindorff-Larsen

## Abstract

Time-resolved experiments can provide unique insights into dynamical processes such as protein folding, ligand binding, and many other molecular processes. These experiments are, however, difficult to interpret at the molecular level because they generally report on signals that are averaged over multiple configurational states, and because they often report on processes that are well beyond what can be studied using most simulation methods. Here we describe an approach to use molecular simulations to model and interpret time-resolved experiments. The method, which we term trBME (time-resolved Bayesian/Maximum Entropy), is based on combining a model for the dynamics of the system, with 3D structural models of the protein that can be compared to the experiments. We illustrate the utility of the model using synthetic time-resolved small-angle X-ray scattering data and show how it can be used to extract detailed information from the experimental data on the process of protein unfolding.

## I. INTRODUCTION

The structure and dynamics of biological macromolecules such as proteins plays a key role in how they function and are regulated. Experimentally, dynamical processes may be studied using a wide range of techniques including nuclear magnetic resonance (NMR) spectroscopy, small-angle X-ray scattering (SAXS), cryogenic-sample electron microscopy, and a range of spectroscopic measurements. Alternatively, biomolecular dynamics may be studied using molecular simulation methods, though these are generally limited in the length- and time-scales accessible, and by the accuracy of the force fields that are used. Often experiments and simulations are combined to generated conformational ensembles that represent the dynamical processes that occur in solution [1]. Such integrative modelling of conformational ensembles is aided by developments in experiments, simulations and methods to integrate the two [1–4].

Several distinct approaches exist to use simulations to interpret experiments [3]. Depending on the goal, these methods need to deal with different sources of error and noise [5], such as limited sampling [6], force field inaccuracies [7, 8] and uncertainties that come from calculating experimental observables from conformational ensembles [9]. We and others have previously argued that Bayesian or Maximum-Entropy-based approaches are particular useful to deal with these kind of problems [2, 3, 10]. In all cases, the goal is to use the experimental data to ‘correct’ or modify a conformational ensemble generated by a computational approach. This modification may either done by introducing a bias during the simulations or by introducing it *a posteriori* after the simulation has been performed [2, 3, 10]. Here, we will refer to such methods to integrate experiments and simulations as *reweighting methods*.

Although several different reweighting techniques are available, most of them deal only with equilibrium (static) experimental data [3], where the experimental observable is often treated as a simple average over the conformations sampled at equilibrium. These methods, however, may not be suitable in two distinct scenarios. First, many experimentally observables such as those probed by NMR and FRET, depend both on the conformational ensemble and the rates at which the different conformations interconvert. We have previously termed these time-dependent data and several methods have recently been developed to help combine such equilibrium-dynamics experiments with simulations [11–15].

Another class of experiments is non-equilibrium, time-resolved experiments where the dynamics is often induced by some rapid perturbation such as a change in temperature, pressure, or solution conditions. Such experiments have the promise to provide ‘molecular movies’ of key biological processes and have provided a wealth of insights [16–27]. The method that we describe here, which we term trBME (time-resolved Bayesian/Maximum Entropy), provides a theoretical and practical framework to use simulations to interpret time-resolved experiments.

Before describing trBME, we describe briefly some alternative approaches to integrate simulations and time-resolved experiments. One approach is to calculate the time-dependence explicitly by averaging over multiple independent non-equilibrium simulations [28], or by linking dynamics in long equilibrium simulations with perturbation-experiments [29? –31]. Such approaches may be very powerful because they directly aim to match the experiments and simulations; however they do have some limitations. First, they are limited by the timescales accessible because the simulations operate on the experimental timescales and require averaging over many simulations. Second, if the simulations and experiments do not agree, there is no clear strategy to use the simulations to interpret the experiments. Another approach tackles the time-scale issue by replacing brute-force simulations of the time-resolved experiment, with an intermediate model with which the dynamics can be studied. In one such example, we previously used simulations to generate a free energy landscape and studied the dynamics of a non-equilibrium, temperature-jump experiments by performing Langevin simulations on a one-dimensional free energy landscape generated by atomistic molecular dynamics simulations [32]. Finally, the ability to correct the simulation-derived dynamics has been studied using the maximum caliber principle (MaxCal) [33], which corresponds roughly to the Maximum Entropy principle, but for paths instead of configurations. While examples are sparse, MaxCal has been used to constrain simulations using time-resolved observations [34–36]. MaxCal is a powerful approach, but the practical implementation to bias simulations is difficult. First, experimental data might be sparse, i.e. collected at broadly separated time points, rendering the bias less ineffective in practice. Second, for complex conformational reactions, the timescales covered by the experiments may be difficult to reach even in the presence of experimental constraints. Third, for many types of experiments, calculating the observables during each step of the simulation is computationally inefficient, and hence may substantially slow down the simulation. Finally, in cases where one wants to examine the effects of different data, it may be a disadvantage that new simulations need to be run for each set of experiments.

Here, we introduce trBME that aims to circumvent the problems outlined above when integrating simulations and time-resolved experiments. trBME is based on two observations. First, many time-resolved experiments consist of a, possibly large, series of experiments that are time-locally static. For example, in a time-resolved SAXS (trSAXS) experiment, each time point corresponds to the ‘instantaneous’ average over the configurations present at that time. This means that we can use existing methods that deal with equilibrium dynamics [3, 10] if we can formulate a kinetic model describing the time evolution of the system between the time points. Second, one of the most central pieces of information that is available from the experiments is the timescales of the non-equilibrium dynamics. Thus, we do not need to predict these timescales from simulations as they are available from the experimental data. Roughly, this means that we can use the simulations to generate the conformational ensembles that represent the non-equilibrium dynamics and use the experiments to derive the timescales. A corollary of this observation is that we can use enhanced sampling methods that aid in sampling slower processes, which generally come at the cost of perturbing the kinetics of the system. In this sense, and as explained in more detail below, our approach shifts the focus to generate realistic time-dependent statistical priors rather than modelling the entire dynamics.

Our trBME approach has the advantage of being able to deal with sparse experimental data and experimental timescales that are unreachable directly by MD because it can be readily interfaced with any enhanced sampling method able to estimate free energy surfaces along a given set of collective variables. Our approach is applied *a posteriori* to running the trajectory, so the computational cost of calculating observables is substantially minimised. We begin by presenting the theoretical and practical framework for trBME and demonstrate it using synthetic data generated to mimic the process of protein unfolding as probed by a temperature-jump, trSAXS experiment.

## II. METHODS

### A. Bayesian Maximum Entropy

As trBME is based on the Bayesian/Maximum Entropy (BME) method for equilibrium data, we briefly present key aspects and refer the reader to previous literature for more details [3, 10, 37, 38]. Given a molecular simulation, whose resulting trajectory {*X*_*i*_}_*i*=1,…,*N*_ is composed by *N* snapshots, we associate a set of weights to this trajectory, 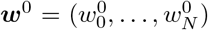 whose *i*-th element reflects the probability to realise the *i*-th configuration given the underlying force field. We refer to ***w***^0^ as the (normalised) *prior*. In the case where extensive sampling is reached by means of an unbiased molecular dynamics or Monte Carlo simulation, the weights are uniform, 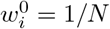. If we instead generate the ensemble using one of many enhanced sampling methods the prior may take less trivial forms that nonetheless often can be estimated [39?, 40].

Given the prior weights, ***w***^0^, one can calculate a set of *M* static observables {*O*_*j*_}_*j*=1,…,*M*_ from the simulation by a weighted ensemble average:

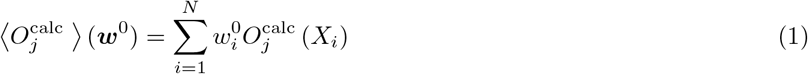

Here 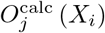 is the value of the *j*-th observable as computed from the *i*-th frame. If the computed observables deviate substantially from the experimentally measured values, *O*^exp^, one may instead use a set of optimised weights, instead of ***w***^0^, in Eq. 1, and optimise those weights to minimise the deviation to experiments. Different options exist to do so [2, 3, 10], but here we focus on the so-called Bayesian Maximum Entropy (BME) method [10, 37, 38]. This approach employs Bayes’ theorem to define a functional *T* (***w***|***w***^0^), which can be minimised to find the optimal set of weights, ***w***^∗^ = min_***w***_ *T* (***w***|***w***^0^), or *posterior* :

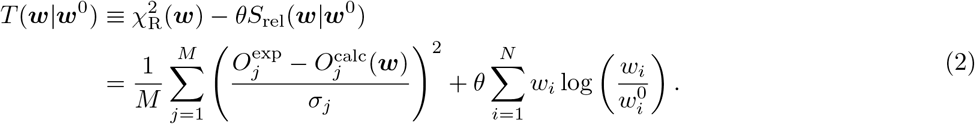

The first term in Eq. 2 represents the reduced *𝓍*^2^ between the experimental and the simulated observables, while the second is the relative entropy *S*_rel_, defined as the negative Kullback-Leibler divergence [41], between the prior and the modified weights ***w***. The entropy term acts to regularise the problem and prevent overfitting of the *M* weights to the *N* observables in *𝓍*^2^, as typically *M >> N*. Finally, *θ* is a positive quantity reflecting our trust in the prior; *θ* = 0 means no weight is put on the prior, while as *θ* increases less and less weight is put on the data. *θ* is usually treated as a hyperparameter, and several methods have used to estimate *θ* [38, 42? –50]. We refer the reader to the Supporting Information for details of how we selected *θ*. We quantify the amount of reweighting using *ϕ*_eff_ = exp(*S*_rel_), which represents the amount of the input simulation that is retained after reweighting [3], though is easiest to interpret when |***w***^0^ is uniform.

### B. Dynamical priors

In Bayesian inference, selecting the right prior can be key to obtaining a meaningful posterior [51]. For static experimental observations, the generation of a meaningful prior might be technically and practically complex but remains conceptually easy to understand, as it only requires assigning meaningful weights to each frame of a given molecular simulation. In the case of time-resolved data, the same *static prior* ***w***^0^ should not simply be applied to each experimental time point, as this would not reflect our knowledge of the time-evolution of the system.

As discussed above, many time-resolved experiments are conducted in quasi-static conditions, making them effectively instantaneous. Therefore, each experimental time point can be considered to represent an ensemble in local equilibrium. While Eq. 1 strictly holds true only for static measurements, we will make use of it in the following to model averages computed for a given experimental time point. This observation enables us to model each time point independently from each other once we obtain a time-dependent prior. For each time point *t*, we would thus have 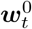.

Furthermore we use two often utilised approximations that enable us to build a set of multiple *dynamical priors* from the same molecular simulation, reflecting our knowledge of the time-dependent behaviour of the system. First, we assume that the dynamics of the system can be approximated using a small set of collective variables (CVs) {*q*_*k*_} _*k*=1,…,*K*_. Second, we assume that the microscopic dynamics in this space can be approximated by an overdamped Langevin equation:

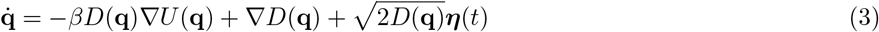

Here *D*(**q**) is a position dependent diffusion coefficient, 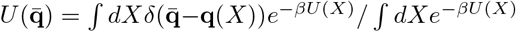 is the potential of mean force, and *β* = (*k*_B_*T*)^−1^. ***η***(*t*) is a Gaussian random noise with ⟨***η***(*t*) ⟩ = 0 and ⟨***η***(*t*), ***η***(*t*^*′*^) ⟩ = *δ*(*t* − *t*^*′*^), where *δ* (·) is the Dirac delta function. The term ∇*D*(**q**) is a ‘spurious drift’ term from following the Itô convention [52].

If the second approximation holds true, the conditional probability density *p* (**q**, *t* | **q**_0_, *t*_0_) for the system to be found in **q** at a given time *t*, provided that we started in **q**_0_ at time *t*_0_ is described by an appropriate Fokker-Planck equation.

In this case it is the Smoluchowski equation:

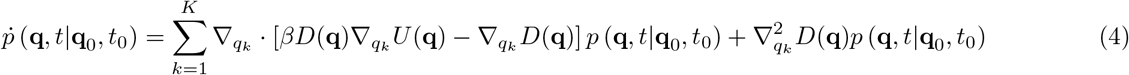

The solution of Eq. 4 enables us to generate multiple time-dependent probability densities that are coherent with our computational knowledge about the time evolution of the system as equilibrium is approached. As long as the initial and the final time points in the experiment are known (e.g. the folded and unfolded state of a protein), it is automatic to convert such probability densities *p* (**q**, *t* | **q**_0_, *t*_0_) into weights 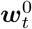 to be used as priors for BME reweighting.

While *D*(**q**) can be estimated both from long trajectories of unbiased simulations and enhanced sampling [53, 54], or estimated from short unbiased trajectories [55], in the present paper we further simplify the calculation, by assuming the diffusion coefficient to be constant *D*(**q**) = *D*, so that its contribution only amounts to an overall rescaling of time 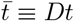 in Eq. 4:

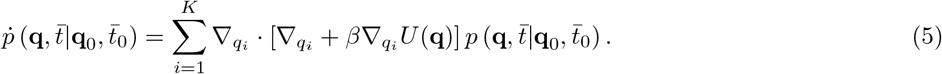

After finding solutions of Eq. 5 we calculate a number of probability densities which is much larger than the number of experimental time points and select the priors based on which densities provide weights optimally fitting the experiments. This also allows us to avoid the dependence on the specific starting point in the simulation of the FP equation. The technical details of the solution of Eq. 5 and the procedure to generate the dynamical priors are discussed in the SI section C. We note that even when the assumptions for the prior do not hold strictly, it ultimately reflects an incomplete prior knowledge of the process under study, and acts as input to refinement using the experiments.

### C. The trBME pipeline

We now have all the ingredients of trBME (Fig. 1). The first steps consist of collecting time-resolved experimental data and running a molecular simulation modelling the processes probed by the experiment. Many different sampling strategies that help define the free energy surface of the system along a given set of collective variables can be employed; these include metadynamics [56], umbrella sampling [57] and parallel tempering [58]. This makes it possible to go beyond the time-scale-limitations of standard MD simulation and increases the chances of sampling the motions relevant for the experiments.

**FIG. 1.**
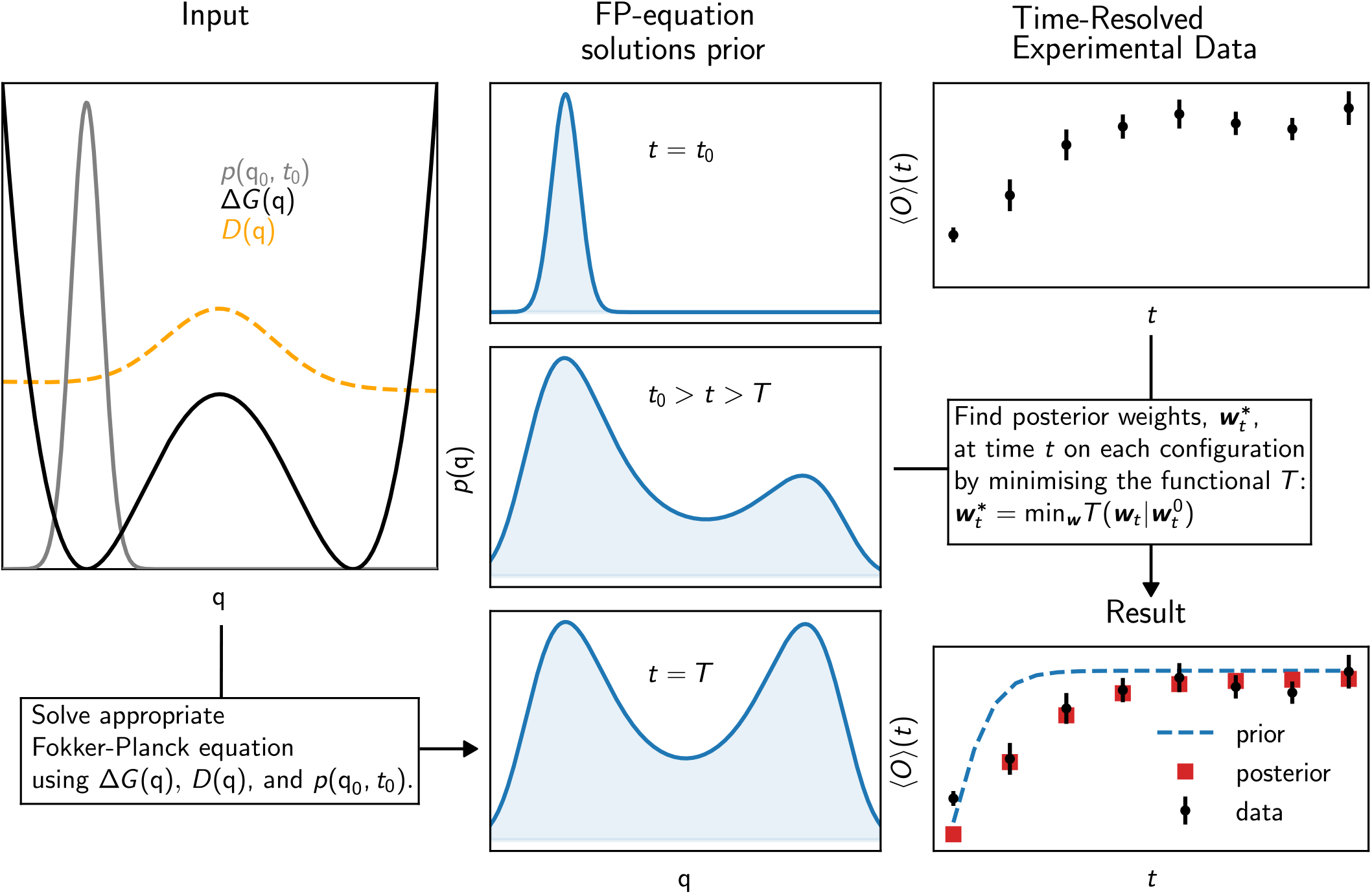
Schematic illustration of the time-resolved Bayesian/Maximum Entropy (trBME) approach to interpret time-resolved experimental data. As input, one needs a molecular simulation that samples configurations of the system together with their corresponding free energies, and —optionally—a diffusion profile along a small set of collective variables *q*. Also an initial distribution of the time-resolved process, *p*(*q*_0_, *t*_0_), is required. Then the free energy, an initial position, and a diffusion profile is used to solve a Fokker-Planck (FP) equation in the space of one or a few collective variables (CVs). This provides a time-dependent probability density from the diffusive dynamics in CV-space. A schematic of this time dependent solution is shown for three different time-points. Equilibrium is reached at *t* = *T*. The time-dependent FP equation solution and time-dependent experimental data, ⟨*O*⟩ (*t*), are combined in a Bayesian/Maximum Entropy fashion. The FP-solution are Bayesian priors and the experiment is data, used to update the priors. Finally, the result of trBME will be time-resolved ensembles of configurations that includes experimental information in a well-defined way. This is illustrated by comparing the experimental observables with simulations before (prior) and after (posterior) reweighting.

In the second step, we simulate the dynamics of the system on the free energy surface obtained from the simulations; Specifically we use this surface as a potential in the Smoluchowski equation and solve this numerically to obtain a series of time-dependent conditional probability densities. These densities are weights over configuration space (parameterised via the CVs) and we use these to calculate averaged experimental observables. We then select the densities that give rise to the best agreement with the experiments. By doing so, we can match the timescale probed by the experiments with that obtained from solving the Smoluchowski equation, giving us what we here term the ‘dynamical prior’.

The third step comprises of the actual time-point-wise reweighting using BME [38]. We treat the experimental measurement at different time points separately, and reweight the computational ensemble using standard BME with the dynamical priors 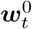.

### D. Temperature-induced unfolding of BSA

In order to demonstrate trBME and test the performance of the method, we generated a synthetic dataset of trSAXS data corresponding to the thermal unfolding of BSA. We selected BSA because it is a relatively large protein (583 amino acids) with a non-trivial folding dynamics. We selected a trSAXS experiment as this method has been employed to probe protein dynamics in a range of systems [16, 19, 22–25], and because we can calculate SAXS data from protein structures accurately and rapidly [59]. We used a C_*α*_-only coarse grained model with a structure-based potential to represent BSA in the simulations [60]. This made it possible directly to simulate longer-timescale motions when generating the synthetic data. As described in more detail below and in the SI, we performed a large number of unbiased simulations at a temperature slightly above the melting temperature of BSA. We added noise to the calculated SAXS data, simulating realistic scattering-angle-dependent noise. Using the same force field and conditions we also ran well-tempered metadynamics (WTMetaD) of the same system, in order to calculate dynamical priors. Details regarding the simulations and the computation of SAXS intensities can be found in the SI sections A and B.

### E. Data and code availability

The code and data used to generate this work is available on GitHub page at https://github.com/KULL-Centre/_2025_hansen_trBME.

## III. RESULTS AND DISCUSSION

To demonstrate and validate trBME we applied it to synthetic dataset generated to describe the dynamics of thermally-induced unfolding of BSA as probed by trSAXS. We performed 1000 50-ns long simulations of unfolding of BSA (Fig. 2C and D), calculated SAXS intensities from each of these, and generated the (synthetic) ‘experimental’ trSAXS signal by averaging across the simulations and adding experimental noise (Fig. 2E and SI Fig. S1). This generates realistic trSAXS data that represents a complex dynamical process with substantial dynamical heterogeneity, and we will hereafter refer to this as the ‘experiments’ and ‘experimental data’. Note that since we use a coarse-grained representation, the simulation time does not represent a physical time, and the real unfolding process would be much longer.

**FIG. 2.**
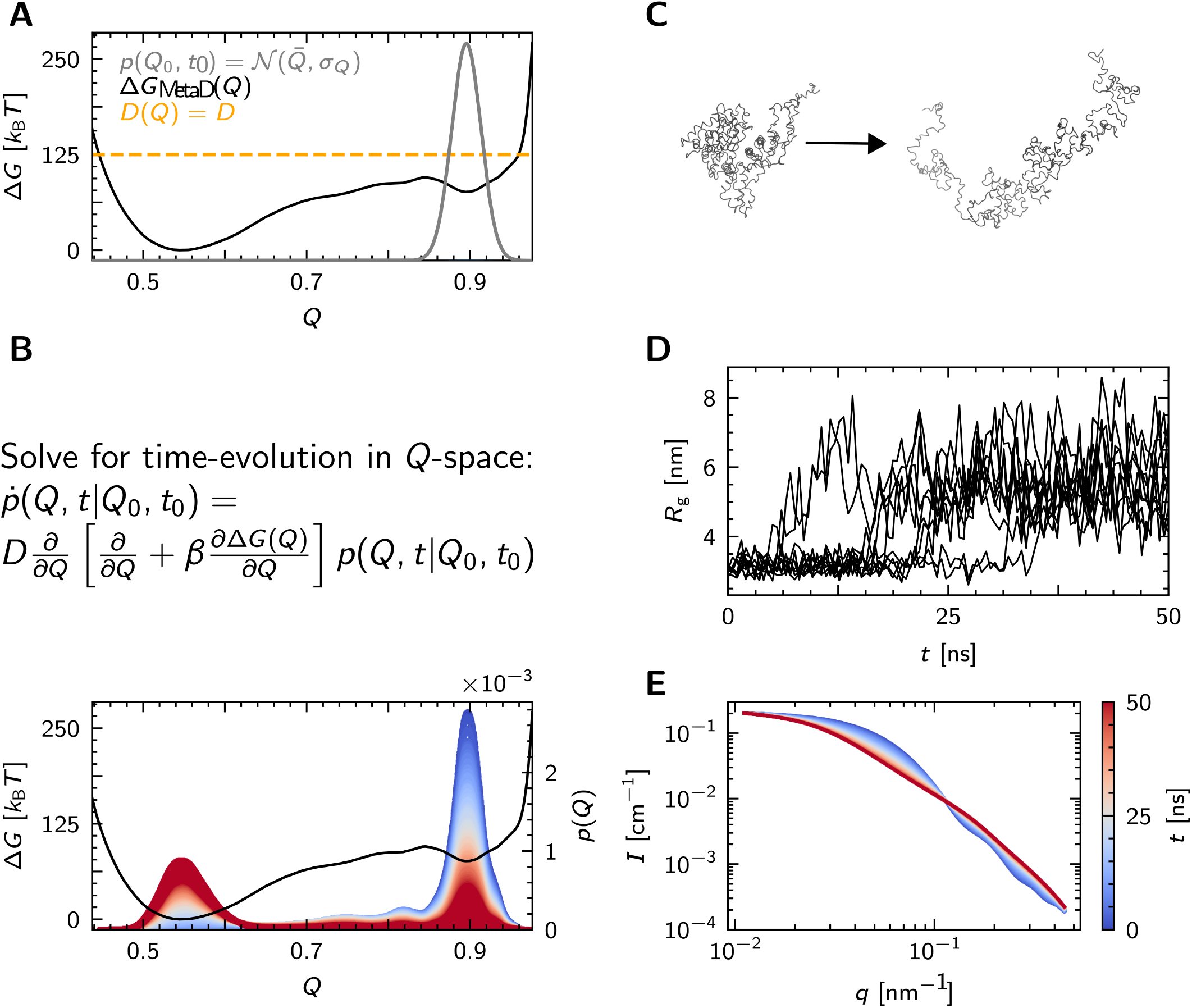
Application of trBME to study unfolding of the protein BSA. **A** The input for trBME from a simulation: A Free energy profile (Δ*G*_MetaD_) generated by a WTMetaD simulation with *Q* the fraction of native contacts as a progress variable, a constant diffusion coefficient, *D*(*Q*) = *D*, and an initial distribution, *p*(*Q*_0_, *t*_0_) set as a Gaussian around the folded basin. **B** The appropriate Fokker-Planck Equation (Smulochowsky Equation) takes the *D*(*Q*), Δ*G*(*Q*), and *p*(*Q*_0_, *t*_0_) as input, and is solved numerically to generate a series of probability density functions, from the initial state to equilibrium. These are shown, coloured by the elapsed time following the colourbar in **E**, alongside the free energy profile. **C** Unbiased simulations of thermally induced unfolding of a C_*α*_ structure-based model of BSA were used to generate synthetic experimental data. Two representative structures showing a partially folded and unfolded state are shown. **D** *R*_g_ over time for unbiased simulations simulations. Shown are 100 of 1000 total unbiased simulations. **E** Synthetic time-resolved SAXS intensities, *I*, as a function of magnitude of the scattering vector, *q*. (We here follow the conventions of naming this SAXS variable *q* and the fraction of native contacts *Q*). All curves are generated from an ensemble average of curves calculated from the unbiased simulations. The curves here are shown without synthetic statistical errors, the curves withs statistical errors are shown in SI section A.

As the input to trBME, we sampled the free energy landscape using a 500-ns-long WTMetaD simulation of the system at the same temperature as that used to generate the synthetic trSAXS data (Fig. 2A). In these simulations we enhance sampling of folding and unfolding of BSA by biasing the fraction of native contacts (*Q*, see the SI section B for details about the simulations). In these simulations we are thus able to observe reversible transitions between the folded and unfolded states and converge to a free energy landscape with multiple minima (Fig. 2A), representing the folded state (*Q* ~ 0.9), the unfolded state (*Q* ~ 0.5) and an intermediate state, spanning *Q* ~ 0.65 − 0.83 (Fig. 2A,B).

The WTMetaD simulations enable us to capture the conformational ensemble and free energy landscape of BSA unfolding (Fig. 2A), but does so in a way that does not preserve the dynamics observed in the experiments. In trBME we re-introduce the dynamics via the generation of ‘dynamical priors’ which are conformational distributions that represent the evolving system at different time points. We generate these by solving Eq. 5 with *Q* collective variable (Fig. 2B), saving distributions at 5000 time points, and selecting the weights that show the lowest 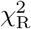 against the experimental data. This an optional procedure, and given a realistic diffusion profile, could be skipped. In Fig. 2B, the corresponding probability densities show how the distribution of conformations moves from the native state towards the unfolded state, and that one or more intermediates are transiently populated during the unfolding process.

Having generated the dynamical priors from the equilibrium simulation, we use these as input to BME to reweight against the trSAXS experiments on time point at a time. We find that using the dynamical priors results in a description that reproduces the experiments well, and find very good agreement between the trSAXS experiments and the results predicted from solving the Smoluchowski equation even before reweighting (Fig. 3 and Fig. S2). One should keep in mind that in this case we optimise the selected timepoints based on 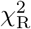, as described above, but is is an optional part of the procedure. The agreement with experiments can be made even better by reweighting, resulting in an averaged 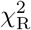 of 1.27±0.02 and an average *ϕ*_eff_ of 0.913±0.004. The high value of the average *ϕ*_eff_ indicates that, as expected, the priors already provide an accurate description of the experiments and that only a small amount of reweighting is required to fit the experiments; we note here that this result is in part due to the fact that the time-point of the prior has been selected using the experimental data.

**FIG. 3.**
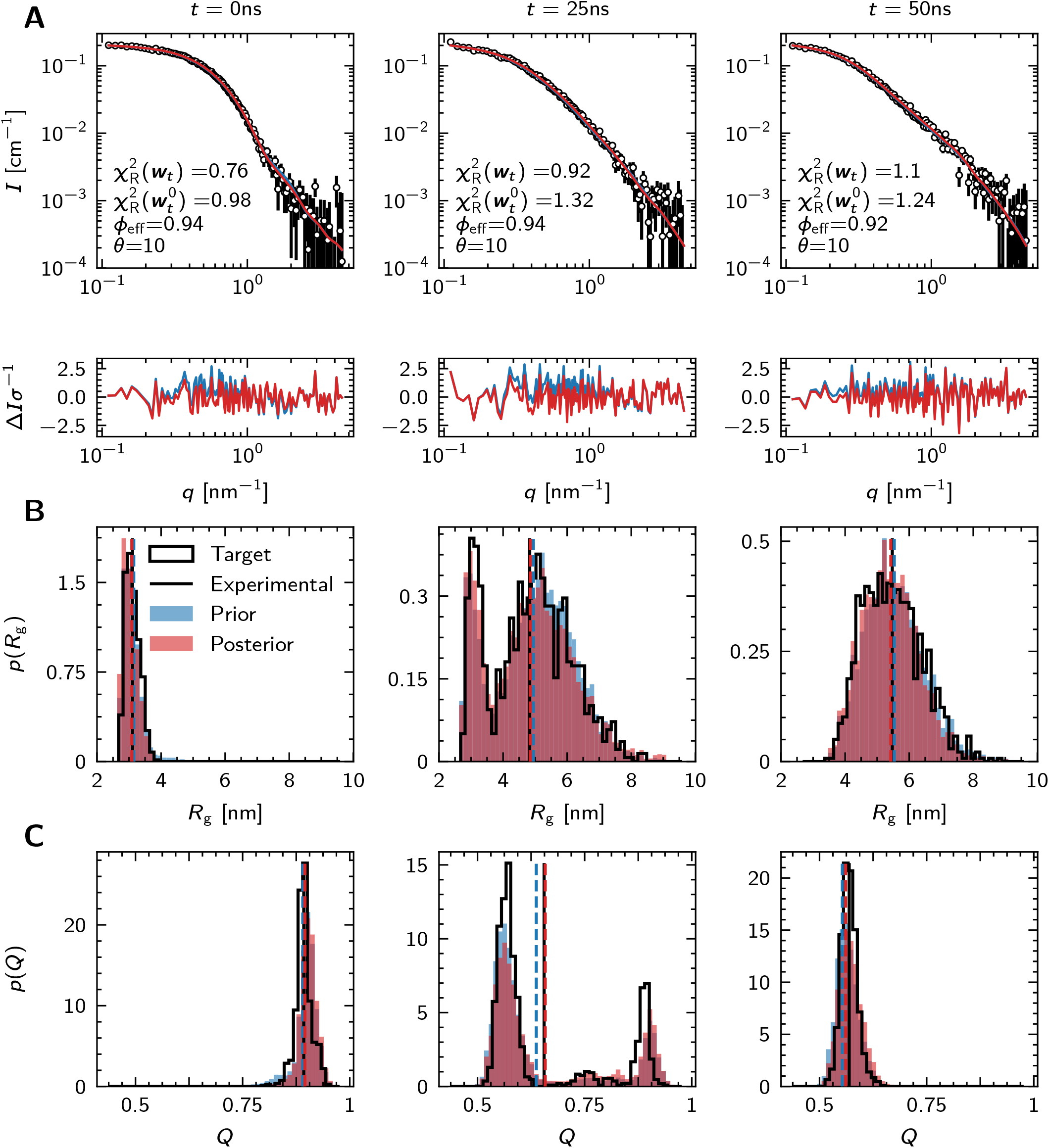
trBME performance visualised at the beginning, midpoint and end of the thermal unfolding of BSA. **A** SAXS curves of synthetic experimental data, SAXS curves generated from ensembles representing dynamical priors and reweighted posterior at time points of 0 ns, 25 ns and 50 ns. We also shown the residuals at each time-point. **B** Distributions of the radius of gyration, *R*_g_. The black outlined histograms correspond to the unbiased simulations used to generate the synthetic experimental data, the blue histograms corresponds to the prior generated from 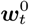, and the red histograms correspond to the results of trBME after reweighting against the SAXS data. Black vertical lines represent the *R*_g_ of the averaged experimental SAXS data and blue and red vertical lines represent the average *R*_*g*_ of the prior and posterior ensembles, respectively. **C** The same as **B** but with *Q* (the fraction of native contacts) as a collective variable.

One advantage of using synthetic data is that they allow us to compare the reweighted ensemble against the time-evolution of conformational ensembles that were used to generate the synthetic data. For example, we compared the probability distributions of the radius of gyration, *R*_*g*_ (Fig. 3B), the fraction of native contacts, *Q* (Fig. 3C), and the RMSD (SI Fig. S3) for several time points. We find that—in addition to matching the experimental observables (which we reweighted against)—the reweighted ensembles match these other properties very well. Together, these results show that the combination of the dynamical priors generated by the Fokker-Planck modelling and reweigting against the experimental data results in conformational ensembles that match the ground truth beyond ensemble-averaged quantities.

To examine how much the success of trBME depends on the dynamical priors that we have introduced and generated, we examined two alternative choices for static (time-independent) priors: the weights corresponding to the bias in the WTMetaD simulation describing the equilibrium ensemble, 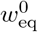, and uniform weights (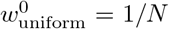 across all *N* frames). We find that while BME reweighting results in reasonable fits to the trSAXS data (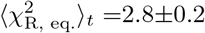 and 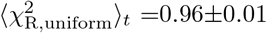 see. Fig. S5 and Fig. S9) and reasonable estimates of the averages of relevant observables, the underlying probability distributions differ substantially from the ground truth used to generate the synthetic data (Fig. S6, S10). This demonstrates that an uninformed choice of priors leads to a less accurate description of the dynamics of the unfolding process, and requires more extensive reweighting to fit the experiments (as seen also by ⟨*ϕ*_eff,eq._⟩ _*t*_ =0.31 ± 0.04 and ⟨ *ϕ*_eff,uniform_ ⟩ _*t*_ =0.77 ± 0.02, which are both substantially lower than ⟨ *ϕ*_eff_ ⟩ _*t*_ of 0.913 ± 0.004 for the dynamical priors) (see SI Fig. S8, Fig. S4, and Fig. S2).

We illustrate these findings by showing the progression of the populations three different states (Fig. 4ABC) in the trBME analysis with the different priors. The definition of the populations are described in SI section F. We find that the populations obtained from trBME with the dynamical priors match very closely (within 5%) those in the original simulations used to generate the SAXS data (compare black and orange lines); in contrast trBME with both the uniform and equilibrium priors give less accurate results for the populations of the three states. This can also be seen by calculating the Jensen-Shannon divergence between the distribution of *Q* from the ground truth and those obtained with trBME with the different priors (Fig. 4D).

**FIG. 4.**
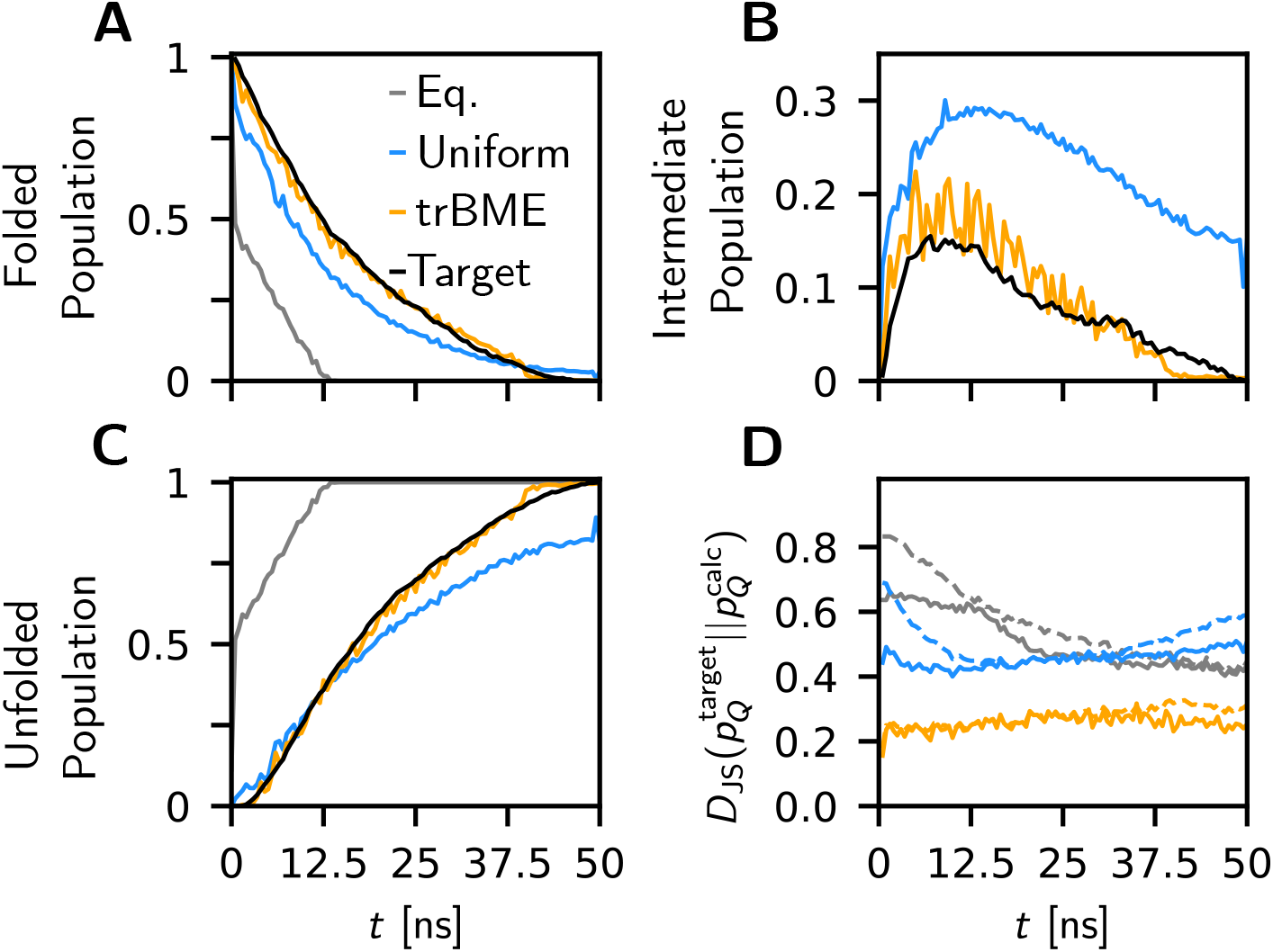
Analysing the effects of using different priors in trBME as a function of synthetic experimental elapsed time. We compared the posteriors of using the dynamical priors from solving the Smoluchowski equation with the results from two different choices of static priors (Equilibrium and Uniform). **A, B, C** Calculated time-evolution of the population of the native, intermediate and unfolded states using different priors (Uniform, Equilibrium and trBME) along with ‘Target’ indicating time-dependent populations from unbiased MD. **D** The Jensen-Shannon divergence between a given ensemble and the corresponding ensemble from unbiased MD. Both distributions are taken along Q, as a function of elapsed time. Dashed lines indicate priors and full lines indicate posteriors. A Jensen-Shannon divergence of zero indicates no divergence to MD ensemble, i.e. perfect agreement.

## IV. CONCLUSIONS

We have described trBME (time-resolved Bayesian/Maximum Entropy) refinement, an approach to use molecular simulations to interpret time-resolved experimental data. trBME is based on two ideas: (i) many time-resolved experiments are time-locally static and (ii) we can use the experiments to obtain the timescales of the non-equilibrium dynamics, and thus use the simulations to interpret the structural ensembles underlying this dynamics. This idea differs from previous work which have often used the simulations to predict the kinetics of the system. Instead, we posit that experiments generally provide relatively easy access to the overall kinetics and timescales, and we instead focus on using the simulations to provide structural interpretations of time-resolved experiments. We have demonstrated trBME using synthetic trSAXS experiments and show excellent agreement against a number of conformational properties. For example, we show how the method is able to capture the time evolution of the population of the different conformational states. We note, however, that trBME is general and can be used to study other types of time-resolved experiments and processes other than protein folding. Also, while we have here used metadynamics simulations to capture the motions probed by experiments, trBME is compatible with other sampling methods and can be extended to use, for example, Markov state models for the underlying dynamics. We envisage that trBME can be used to provide new insights into the dynamical motions of macromolecules that underlie biological functions.

## ACKNOWLEDGMENTS

We acknowledge support from the NordForsk Nordic Neutron Science Programme (grant number 81912) and the Lundbeck Foundation BRAINSTRUC structural biology initiative (R155-2015-2666). The work was co-funded by the European Union (ERC, DynaPLIX, SyG-2022 101071843). Views and opinions expressed are however those of the authors only and do not necessarily reflect those of the European Union or the European Research Council. Neither the European Union nor the granting authority can be held responsible for them. We also acknowledge access to computational resources from the ROBUST Resource for Biomolecular Simulations (supported by the Novo Nordisk Foundation grant no. NF18OC0032608), the Danish National Supercomputer for Life Sciences (Computerome) and the Biocomputing Core Facility at the Department of Biology, University of Copenhagen.

## Appendix A: Generating reference time-resolved ensembles and synthetic SAXS data

### 1. Molecular dynamics simulations

We built a structure-based C-*α* model using the SMOG web-server [61], where the contacts were defined through a shadow map [62]. In order to avoid self-interactions with periodic images in the unfolded state, we built a box with 5 nm padding between the protein and the box limits. Each *C*_*α*_ bead was assigned with unit mass and zero charge. Finally, we manually modified the force field to represent the native protein’s disulphide bonds as bonds. In particular, covalent bonds were added between the following residue pairs: (53,62), (75,91), (90,101), (123,168), (167,176), (199,245), (244,252), (264,278), (277,288), (315,360), (359,368), (391,437), (436,447), (460,476), (475,486), (513,558), and (566,557). In this way, our simulations represent the unfolding of oxidised BSA.

To simulate a temperature-jump unfolding experiment, we started multiple MD simulations (using GROMACS v.2019.6) from a native state crystal structure (PDB: 4F5S, [63]). The equations of motion were integrated using a Langevin integrator with 0.5 fs timestep. We performed 50-ns-long simulations at a temperature of 161 K, which is just above the melting temperature of the structure-based model (see main text section B). From these initial simulations, we selected 1000 trajectories where we observed unfolding within the 50 ns simulation time, and sampled 100 equally-spaced frames from each simulation.

### 2. SAXS curve generation

To generate SAXS curves from the C_*α*_-only structures, we wanted to include contributions from the side chains to the scattering profile. We thus constructed all-atom structures from the C_*α*_ models using PULCHRA v.3.06 [64] before calculating the SAXS curves.

We used PEPSI-SAXS v.2.4 [59] to calculate SAXS profiles. Since our goal was to use it as a forward model rather than a SAXS fitting tool, we fixed the free parameters in PEPSI-SAXS. To do so we we fitted a reference BSA SAXS intensity experiment (SASDE75 [65]) using the BSA crystal structure as an input and consequently determined the three parameters in PEPSI-SAXS as *I*(0) = 0.2 cm^−1^, *r*_0_ = 1.66 Å and *δρ* = 1.6%; these parameters were then fixed in the 10^5^ SAXS calculations (1000 simulations with 100 frames from each). Finally, in order to mimic experimental noise, we re-sampled each intensity point by drawing it from a Gaussian distribution centred on the calculated intensity value, and using the *q*-dependent experimental error from the reference experiment as standard deviation: *I*_noisy_(*q*) ~ 𝒩 (*I*_average_(*q*), *σ*_exp_(*q*)). In Fig. S1, we show the averaged and noisy intensities that we used as input to our reweighting calculations.

## Appendix B: Well-tempered metadynamics simulations

In order to select the simulation temperature to mimic a temperature unfolding simulation, we ran multiple short well-tempered metadynamics simulations at different temperatures with the goal of identifying the melting temperature of BSA in the structure-based model. We selected 12 temperatures in the range 120 K–175 K, and ran metadynamics at each temperature using the fraction of native contacts, Q, as a collective variable for the bias. We defined the collective variable as [66]:

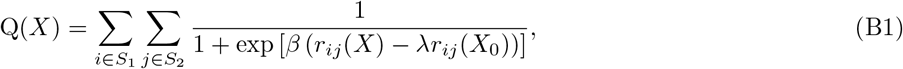

where λ = 1.2 and *β* = 50 nm^−1^. As contacts in the calculations of *Q* we used the same contacts that define the structure-based model, ensuring that the collective variable captures key aspects of the model. In the metadynamics simulations we used a bias factor *γ*^−1^ = 100, a hills height of 4 kJ/mol, *σ*_*Q*_ = 0.01 and a deposition rate of 0.5 ps.

Based on the short metaD simulations at different temperatures, we selected *T*_exp_ =161K as the synthetic experimental temperature for our reference dataset and ran 0.5 *µ*s using the same setup and forcefield described above.

We assessed the convergence of the WTMetaD simulation by means of block error analysis [67]. As shown in Fig. S4 A, the error plateaus between 50 ns–100 ns of block length, and therefore we consider it converged. Provided the convergence of the metadynamics run, we computed the weights for all the frames in our simulation by Torrie-Valleau reweighting [39] using Plumed’s REWEIGHT BIAS function, and then employed these weights as the equilibrium (eq.) prior.

**FIG. S1.**
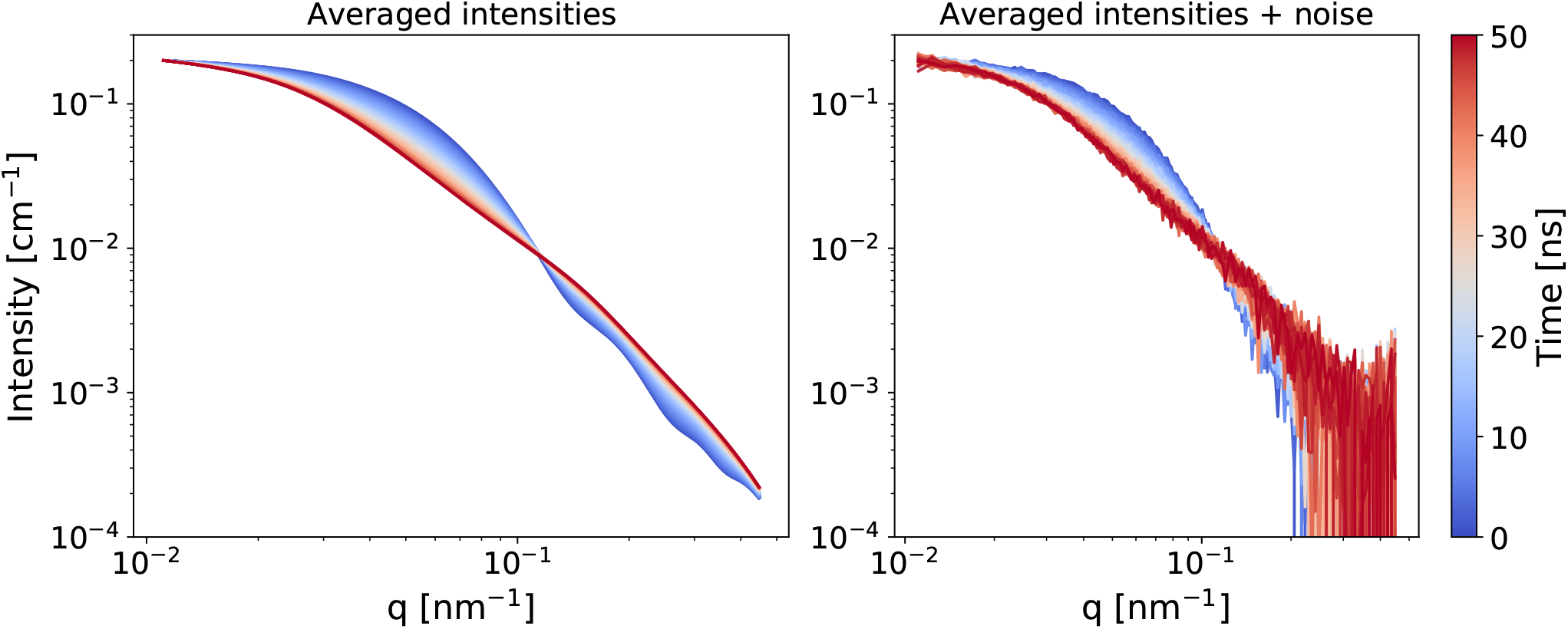
Synthetic trSAXS data for temperature-induced unfolding of BSA. Left: SAXS intensity curves calculated from 1000 unfolding simulations. Right: Same as the plot on the left, but after addition of noise. This is the synthetic data that we used as experimental input to BME. The colours show the time-points (simulation timescale) at which the ensemble-averaged SAXS curve were calculated.

## Appendix C: Dynamical priors

### 1. Construction of dynamical priors

To generate the dynamical priors, we solve a one dimensional Fokker-Planck equation numerically by discretising the spatial and temporal components, and solving with the python package FPlanck [68]. The package is intended for modelling diffusion in physical space, and we therefore have to select parameters with units related to nm and convert back to our CV-units later. The input parameters reported here are in CV-units, as the Fokker-Planck units have no physical relevance. When generating the dynamical prior, we set the initial state, *p*(*Q*_0_, *t*_0_), to a Gaussian distribution centred at 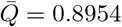, with a *σ*_*Q*_ = 0.0180, approximating the centre and width of the folded basin.

The numerical FP-solver was run with a grid size of 50 for 5000 time points. From these 5000 time points, we selected100 points that minimised the 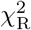 to experiment before reweighting, under the constraint that the time-ordering is preserved. We perform this selection to help ensure that the priors reflect the time-scale in the experiment. In the case where the CV-dependent diffusion coefficient is known, simply finding the point where the process reaches the final experimental measurement could be sufficient. The drag (or diffusion coefficient) was set to a constant arbitrary value, as the rescaling of the frames will make the exact value unimportant (see main text Eq. 5). The free energy calculated from metadynamics was converted to Joule and used as the potential *U*, along with the temperature, *T* = 161K.

### 2. Determination of *θ*

For trBME, we used a ‘knee-locator’ for each time-point to find a point on a (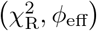, *ϕ*_eff_)-curve to find an optimal *θ*, 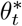. This method has the advantage of being general, but depends on an ‘L’-like shape of a (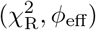, *ϕ*_eff_)-curve. This was done with the Kneed python package (pypi.org/project/kneed/) [69]. For further details, see Bottaro et al. 2020. [38].

For the equilibrium prior we used a constant ***θ*** across all time-points as no ‘L’-shape was visible in the (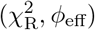, *ϕ*_eff_)-curve (see Fig. S4 and Appendix D).

**FIG. S2.**
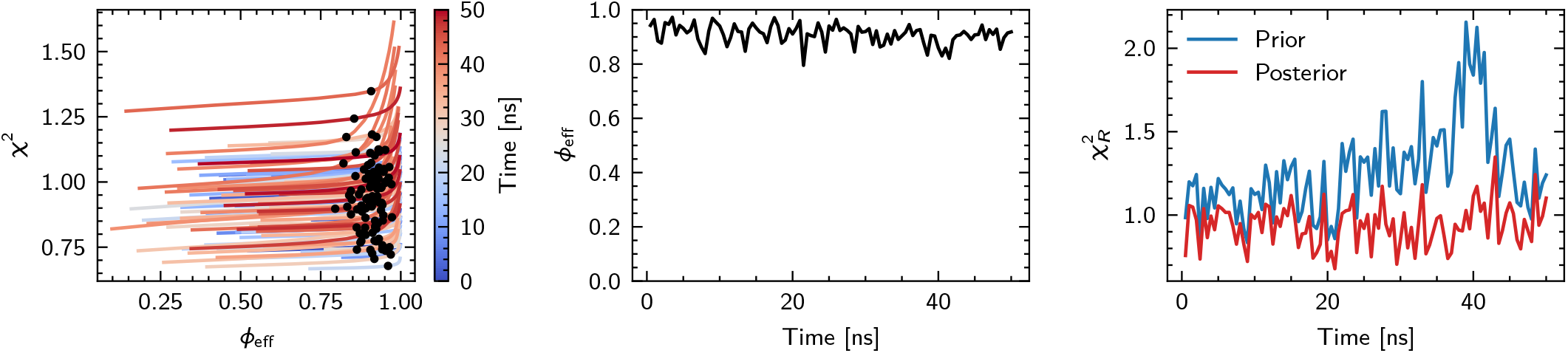
Results with the Dynamical Prior. Left: 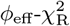-plot. Colour denotes the value of *t*. Black dots indicate 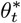 Middle:*ϕ*_eff_ as a function of *t*, after reweighing at 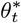. Right: 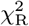, from the SAXS curves, before and after reweighing at 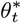. Results for all 100 points are shown here.

**FIG. S3.**
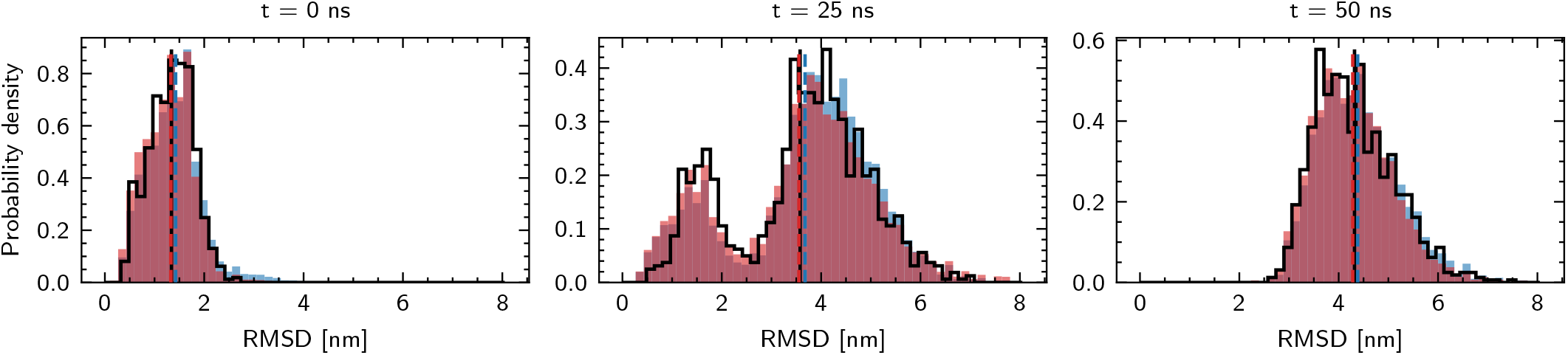
Results for three selected timepoints with the Dynamical Prior using the RMSD from the starting structure as variable. The black outlined histograms correspond to the unbiased simulations used to generate the synthetic experimental data, red histograms correspond to the results of trBME after reweighting against the SAXS data at the optimized *θ*-value, 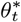, and the blue histograms corresponds to the prior generated from 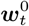. Black vertical lines represent the RMSD of the averaged experimental SAXS data and blue and red vertical lines represent the average RMSD of the prior and posterior ensembles, respectively.

### 3. Results from the Dynamical Prior

For the primary results of the dynamical prior, see main text Fig. 3 and Fig. 4 as well as Fig. S2 and Fig. S3.

## Appendix D: Equilibrium prior

### 1. Equilibrium Prior

In this section we will discuss the results obtained by employing the constant prior provided by the Metadynamics estimate of the equilibrium distribution. In this case, the reweighting is carried out with the same prior, provided by the unbiasing metadynamics weights, for every experimental time point. The *θ* values employed in the reweighting range from 1 to 600. In order to select the optimal value of *θ* for each experimental time point, we looked for an ‘L’-shape in the 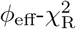 curves, but the majority of these curves did not show a clear ‘L’-shaped decreasing of the 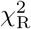, rendering it difficult to associate an optimal *θ* value to our reweighting. Therefore, we selected a constant *θ*^∗^ = 50 for all the frames. The behaviour of *ϕ*_eff_(*t*) (Fig. S4) follows the expectations: it smoothly switches from *ϕ*_eff_(*t* = 0) ~ 0 (as the reference ensemble is represented by fully folded proteins while the prior is, instead, characterised by fully unfolded proteins) to *ϕ*_eff_(*t* = 50) ~ 1 (where instead the prior and the reference ensembles almost exactly match).

A coherent trend is followed by the prior and posterior 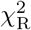 as a function of time: the posterior 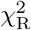 is, as expected, always lower than the prior for every frame and both of them are monotonically decreasing with time, suggesting that it is easier to fit the reference ensemble for long times (i.e. unfolded protein) rather than for short times (i.e. folded protein). Visual inspection of the SAXS fits (Fig. S5) confirms this trend.

The distributions of the radii of gyration and fraction of native contacts obtained from reweighting with the reference distributions are reported in Fig. S6 and Fig. S7, along *Q* and *R*_*g*_ respectively. While the average values are generallycaptured well(less than 5% deviation for Rg and less than 20% deviation for *Q*), the distributions at each time-point do not qualitatively capture the underlying distributions, with the notable exception of the time-points close to 50 ns. At early time-points, only a few configurations have large weights, and the underlying ensemble is not recreated well. This is because a choice of a constant and mostly unfolded prior forces the BME to pick conformations with low *w*_0_ that are folded to match the SAXS curves and therefore also the average radius of gyration, disregarding the underlying shape of the reference distribution. The issue is alleviated as at larger *t* (e.g. from 35 ns onwards) as more of the ensemble is unfolded, in correspondence also of higher values of *ϕ*_eff_.

**FIG. S4.**
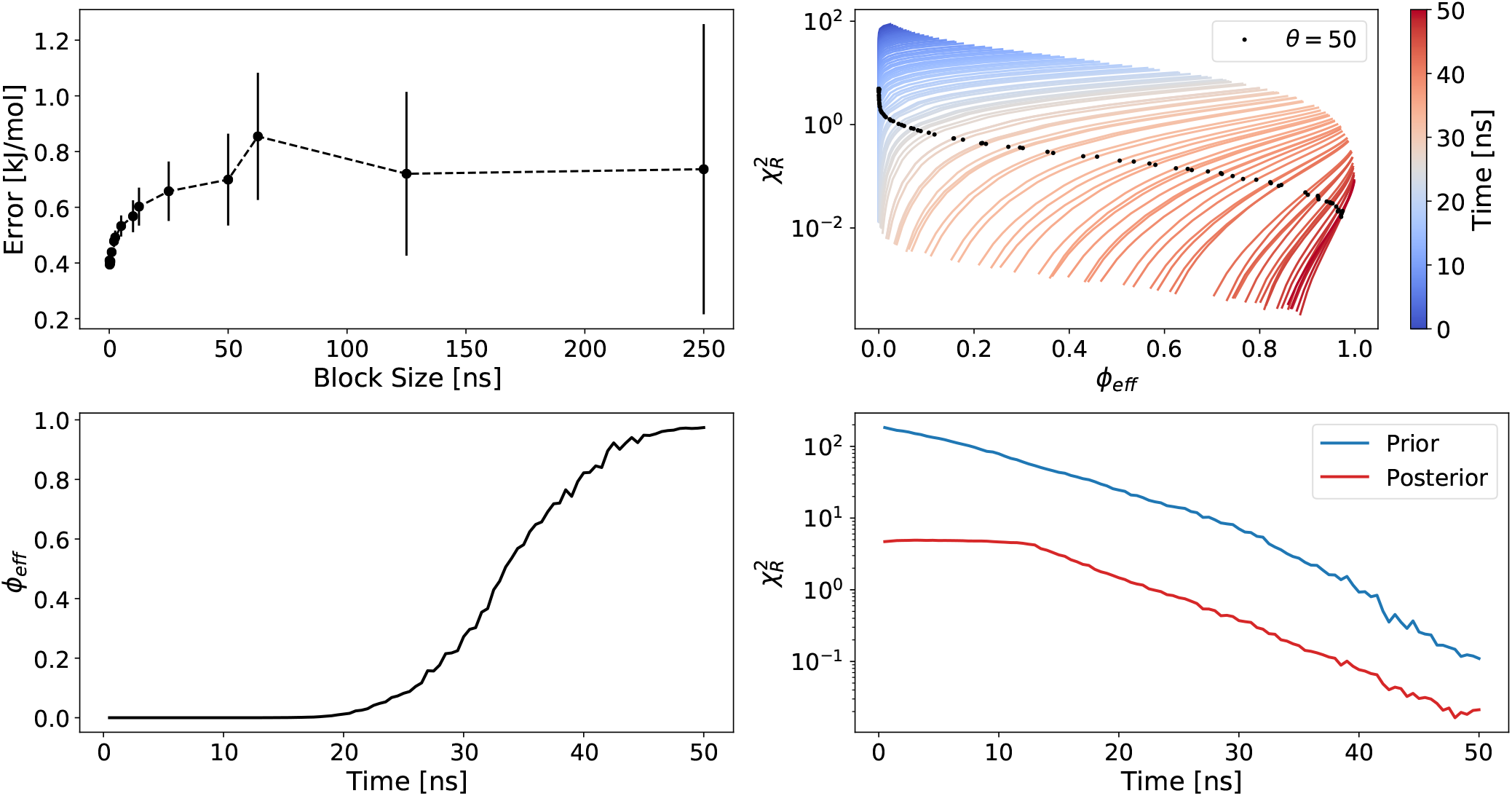
Top left: Error on the free energy, as estimated with block transformations. Top right: 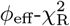 plot. All time points are overlayed. Since no knee-shape is available, a single *θ*-value was selected. This value is shown in black dots. Bottom left: *ϕ*_eff_ as a function of the time-point. Bottom right: 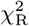 as a function of the time-point. Both prior and posterior is shown.

**FIG. S5.**
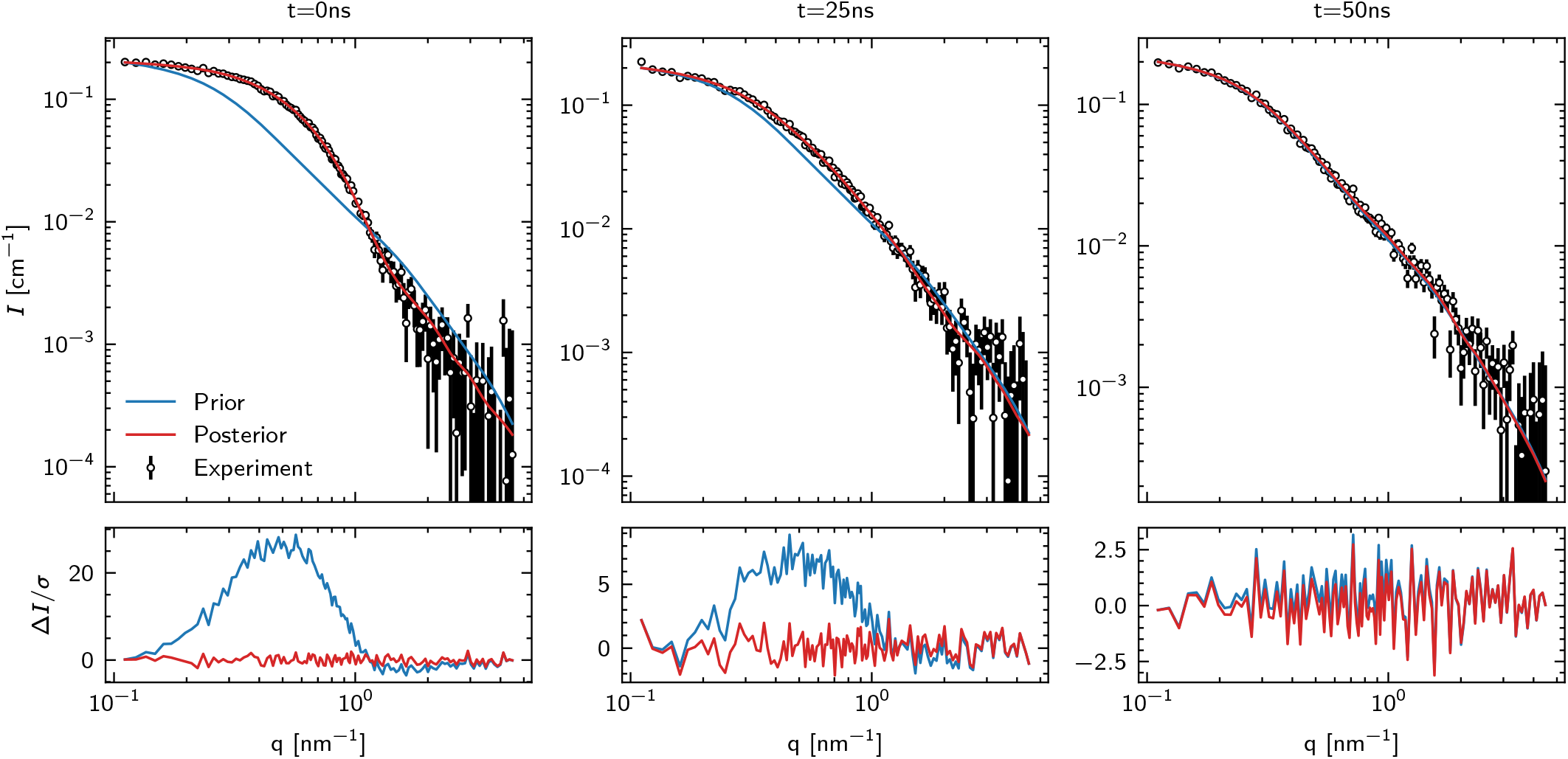
SAXS fits, at three time-points, using the equilibrium prior. Residuals are shown at the bottom, for both the prior and posterior. Posteriors are computed using 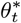.

**FIG. S6.**
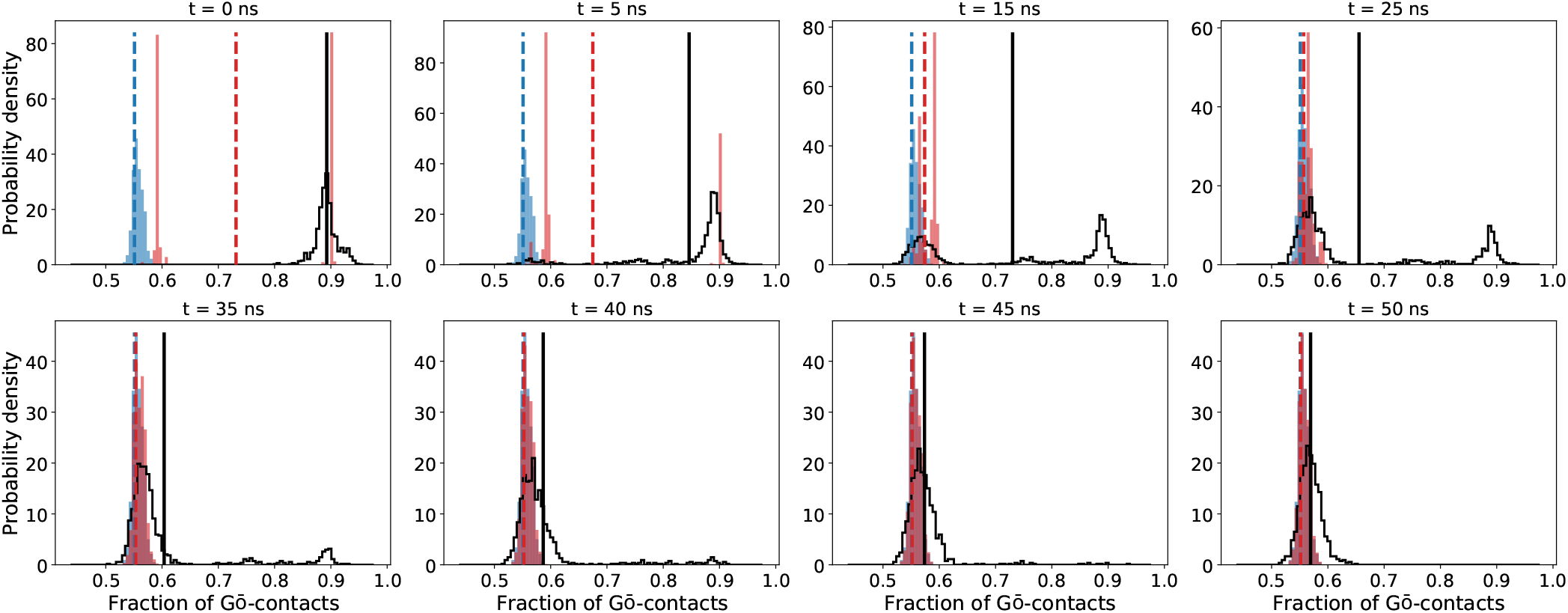
Results of the equilibrium prior, Histograms of *Q* at different time-points. Averages are shown by vertical lines. For histograms and averages, the target is shown as a black outline,the prior and posterior at 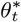 is blue and red, respectively.

**FIG. S7.**
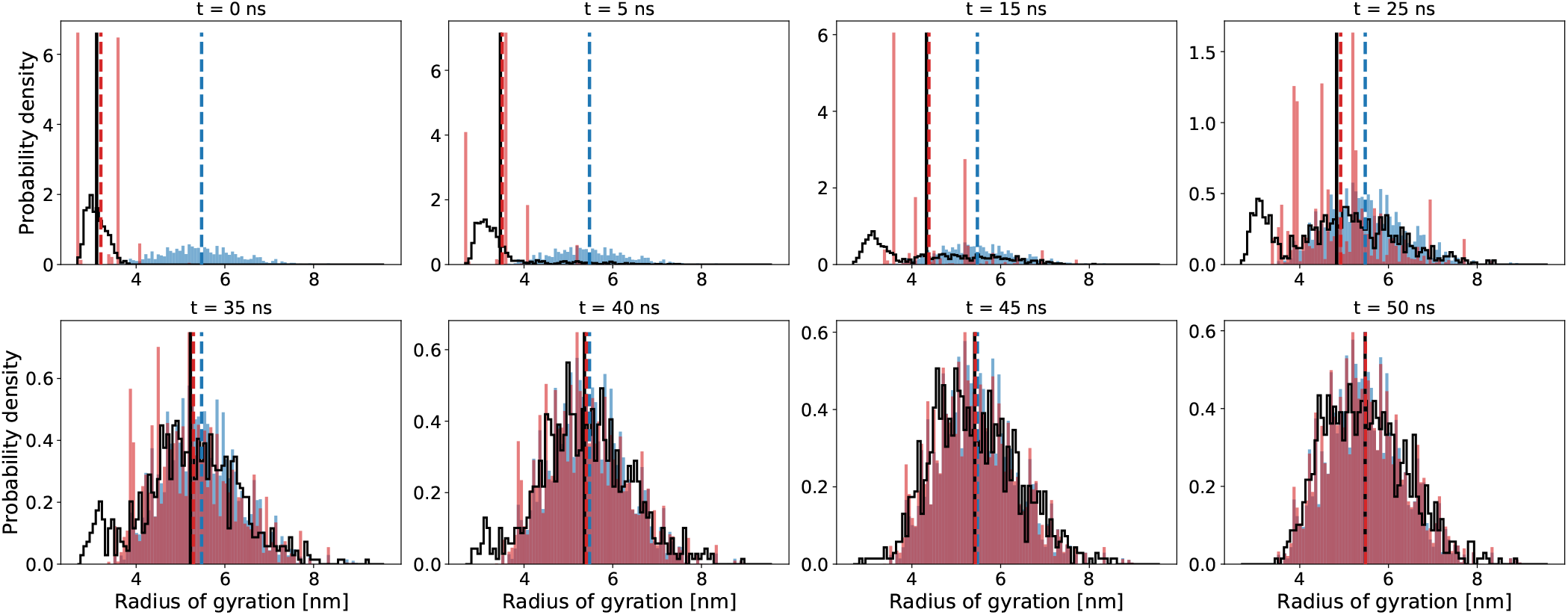
Results of the equilibrium prior, Histograms of *R*_g_ at different time-points. Averages are shown by vertical lines. For histograms and averages, the target is shown as a black outline, the prior and posterior is blue and red, respectively. The bin width is identical for all time-points, and set to highlight the non-smooth nature of the weighted distributions at early time points.

**FIG. S8.**
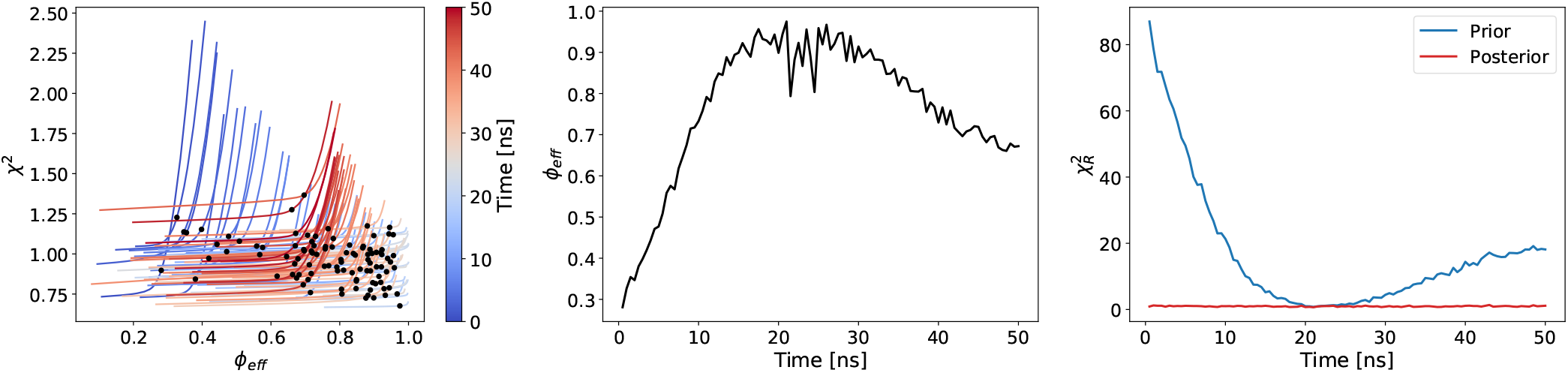
Results of the uniform prior. Left: 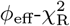-plot. Colours denote the value of *t*. Black dots indicate 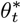 Middle:*ϕ*_eff_ as a function of *t*, after reweighing at 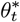. Right: 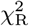, from the SAXS curves, before and after reweighing at 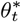.

## Appendix E: Uniform prior

In this section we discuss the results obtained by employing the constant uniform prior provided by associating a constant weight to each frame in the metadynamics simulation, 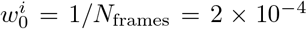, ∀*i*. We stress here that there is no clear theoretical basis to justify this approach, as the metadynamics weights carry information about the bias that has been applied throughout the simulation. Instead, we use the model simply to illustrate the results that one may obtain if the prior does not reflect the time-dependency of the experiments. This prior represents a knowledge of the system where all sampled conformations are equally likely at all time points, and thus could represent a reweighting strategy where a static general prior used, and the experimental data injects dynamic information.

As in the case of the equilibrium prior, the *θ* values employed in the reweighting ranged from 1 to 600. In contrast to the equilibrium prior case, however, it was possible in this case to determine the location of an elbow for all the 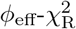 trend is also visible in the prior different experimental time points. The average value of *θ*^∗^ obtained in this way is ≈40. The behaviour of *ϕ*_eff_ as a function of time (Fig. S8) differs substantially from the equilibrium prior, increasing from 0.3 at time *t* = 0 ns and reaching *ϕ*_eff_ 1 around *t* = 25 ns. This is because when the prior is uniform for each configuration, it is most similar to the underlying ground truth when the experiment is probing an ensemble where folded and unfolded configurations are equi-probable. A similar trend is also visible in the prior 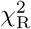, while the posterior 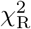 is instead close to 1 for every time point. This is also seen in the SAXS-fits Fig. S9.

For the populations along the radius of gyration (Fig. S11), we see a good agreement with the underlying target distributions for large values of *t*. For low, and intermediate *t*-values, the histograms show a qualitative agreement. The fraction of native contacts is shown in Fig. S10. These show qualitative agreement with the reference ensembles only for the small and large values of *t*. For the intermediate time-points, the posterior is quite flat, and does not reflect the multimodal nature of the target distributions.

## Appendix F: Populations of BSA

The folded, unfolded and intermediate states were determined using two cutoffs along *Q*. The cutoffs were found by fitting a Gaussian to each of four observed peaks and finding their intersection. The cutoffs used where at *c*_*Q*,min_ = 0.675 and *c*_*Q*,max_ = 0.835. The folded population is then all *Q > c*_*Q*,max_, the unfolded is *Q < c*_*Q*,min_, and all others are defined as intermediate.

**FIG. S9.**
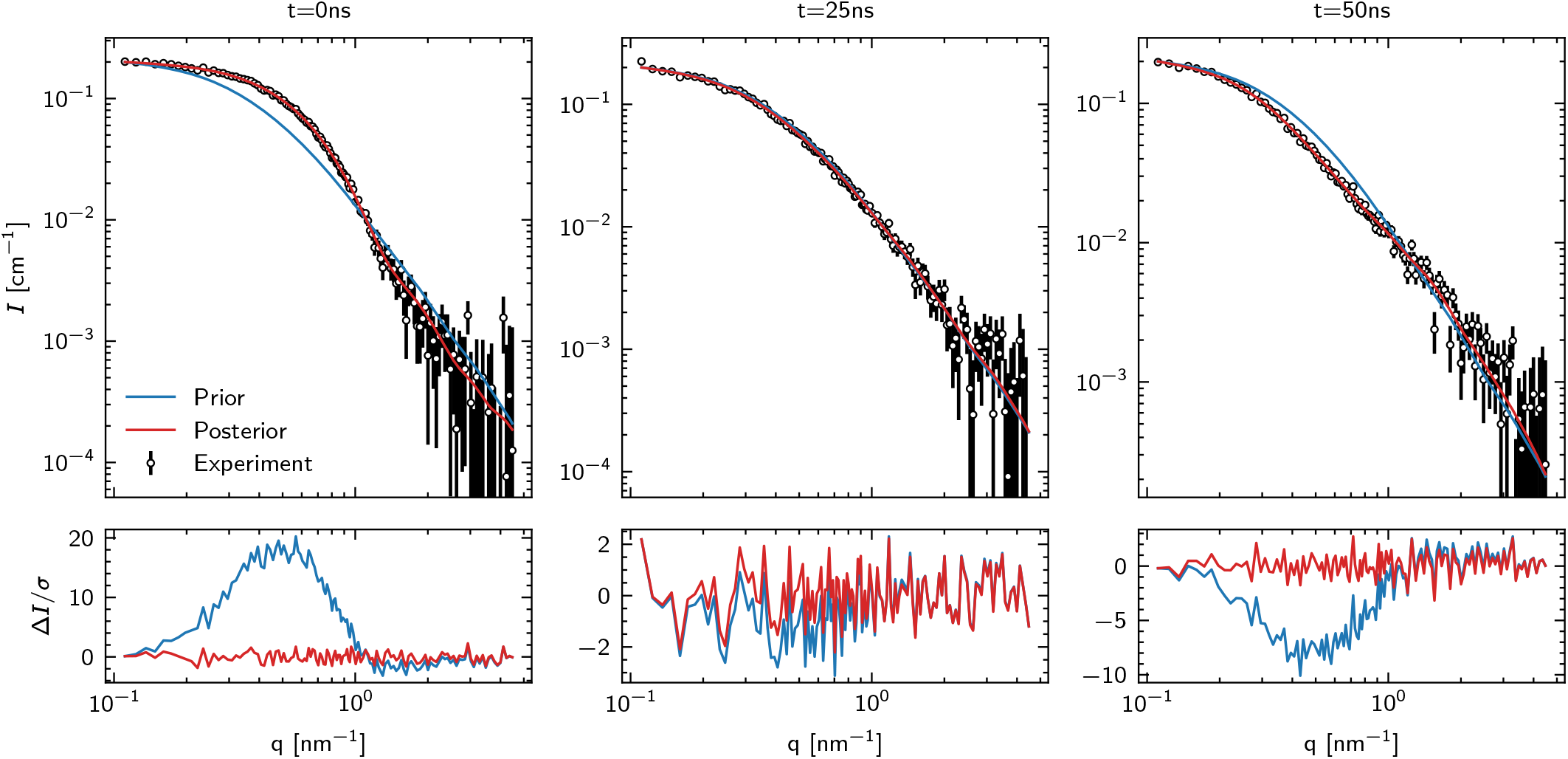
SAXS fits, at three time-points, using the uniform prior. Residuals are shown at the bottom, for both the prior and posterior. Posteriors are computed using 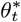.

**FIG. S10.**
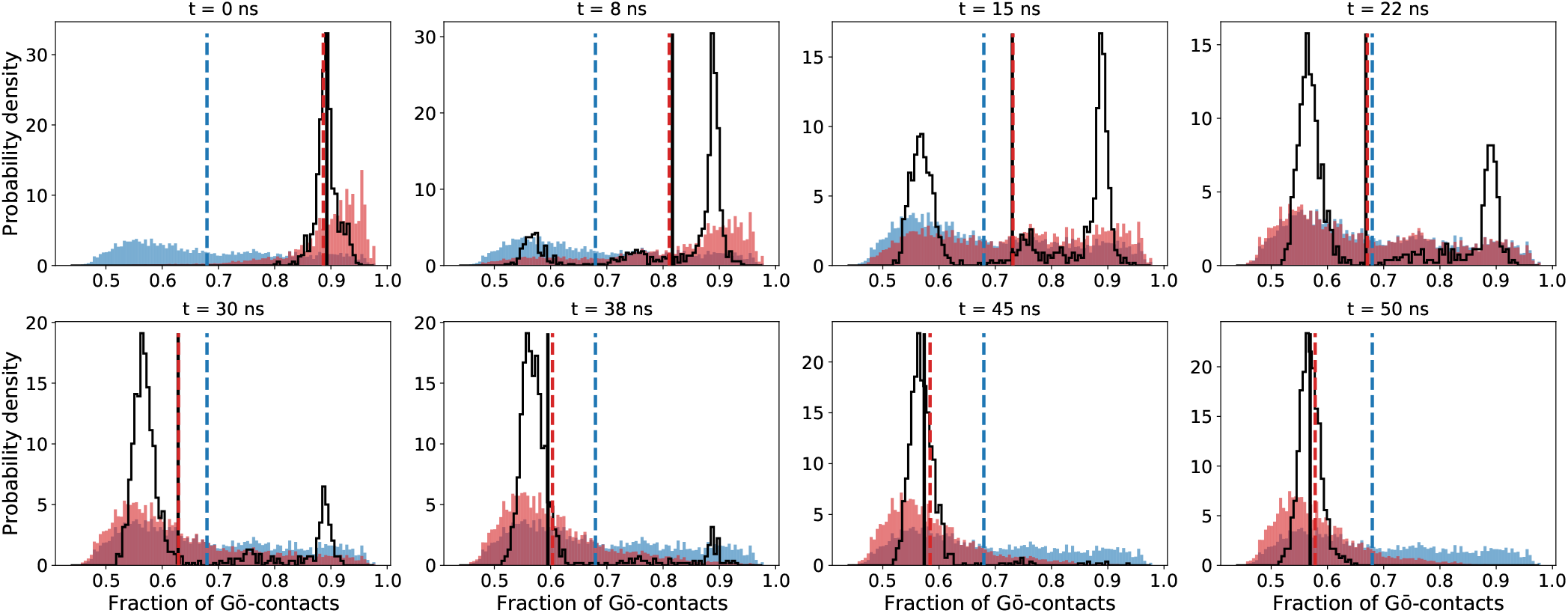
Results of the uniform prior, Histograms of *R*_g_ at different time-points. Averages are shown by vertical lines. For histograms and averages, the target is shown as a black outline, the prior and posterior computed using 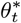. is blue and red, respectively.

**FIG. S11.**
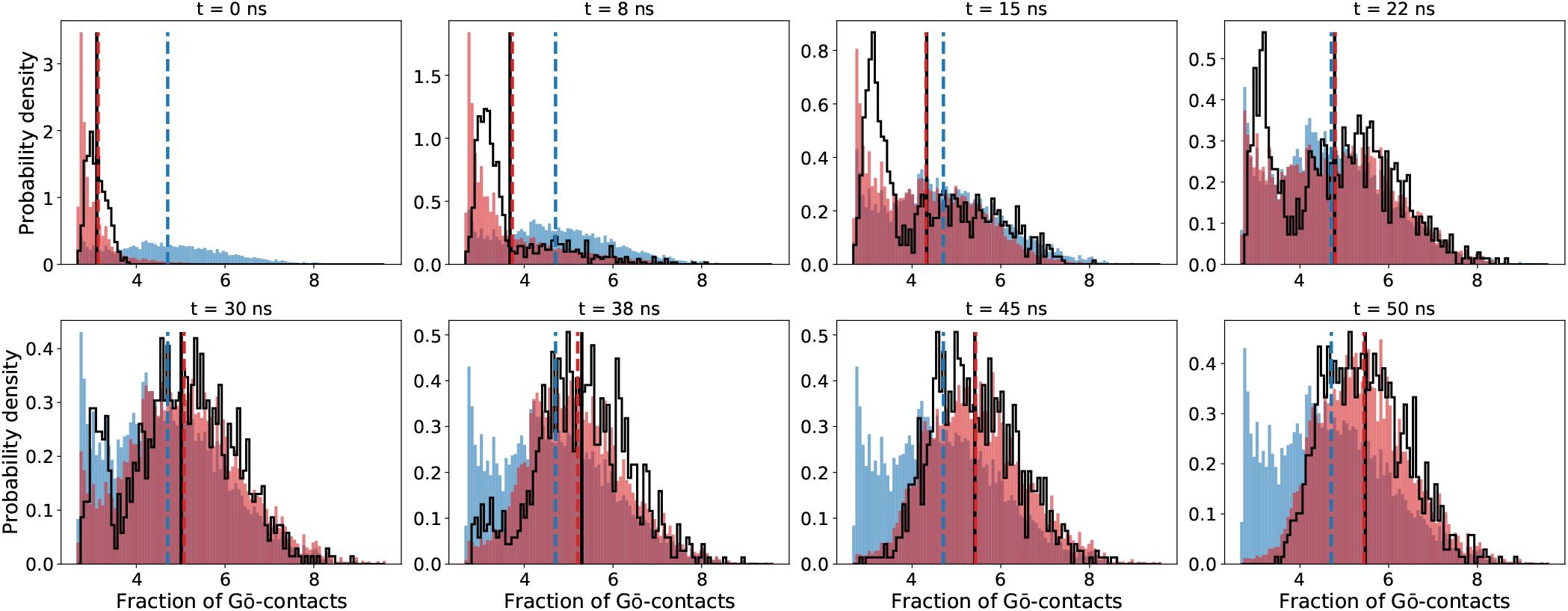
Results of the uniform prior, Histograms of *Q* at different time-points. Averages are shown by vertical lines. For histograms and averages, the target is shown as a black outline, the prior and posterior computed at 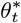 is blue and red, respectively.

